# Global accumulation of circRNAs during aging in *Caenorhabditis elegans*

**DOI:** 10.1101/175026

**Authors:** Mariela Cortés-López, Matthew Gruner, Daphne A. Cooper, Hannah N. Gruner, Alexandru-Ioan Voda, Alexander van der Linden, Pedro Miura

**Affiliations:** University of Nevada, Reno, Department of Biology, 1664 N. Virginia St, Reno, NV, 89557, USA

**Author notes:** corresponding authors: Correspondence to Pedro Miura, Correspondence to Alexander van der Linden. these authors should be considered co-first authors (Mariela Cortés-López, Matthew Gruner).

**Keywords:** circRNA, C. elegans, aging, RNA-seq, splicing, age-accumulation, gene expression

## Abstract

Circular RNAs (CircRNAs) are a newly appreciated class of RNAs that lack free 5´ and 3´ ends, are expressed by the thousands in diverse forms of life, and are mostly of enigmatic function. Ostensibly due to their resistance to exonucleases, circRNAs are known to be exceptionally stable. Here, we examined the global profile of circRNAs in *C. elegans* during aging by performing ribo-depleted total RNA-seq from the fourth larval stage (L4) through 10-day old adults. Using stringent bioinformatic criteria and experimental validation, we annotated 1,166 circRNAs, including 575 newly discovered circRNAs. These circRNAs were derived from 797 genes with diverse functions, including genes involved in the determination of lifespan. A massive accumulation of circRNAs during aging was uncovered. Many hundreds of circRNAs were significantly increased among the aging time-points and increases of select circRNAs by over 40-fold during aging were quantified by qRT-PCR. The age-accumulation of circRNAs was not accompanied by increased expression of linear RNAs from the same host genes. We attribute the global scale of circRNA age-accumulation to the high composition of postmitotic cells in adult *C. elegans*, coupled with the high resistance of circRNAs to decay. These findings suggest that the exceptional stability of circRNAs might explain age-accumulation trends observed from neural tissues of other organisms, which also have a high composition of post-mitotic cells. Given the suitability of *C. elegans* for aging research, it is now poised as an excellent model system to determine if there are functional consequences of circRNA accumulation during aging.

## Introduction

Circular RNAs (circRNAs) have recently been identified as a natural occurring family of widespread and diverse endogenous RNAs [1, 2]. They are highly stable molecules mostly generated by backsplicing events from known protein-coding genes. The expression trends of circRNAs are only recently emerging thanks to RNA-seq library preparation methods that deplete ribosomal RNA (ribo-depletion) rather than enrich for polyadenylated RNA. Most circRNAs are derived from protein-coding genes, and thus one challenge in mapping and quantifying circRNAs is to distinguish reads that can be uniquely ascribed to circular molecules versus linear RNAs emanating from the same gene. Elements located within introns flanking circularizing exons play a role in promoting circRNA biogenesis [3–6], and several RNA binding proteins and splicing factors have been shown to influence circRNA expression [4, 7–10].

Despite the recent attention on circRNAs, their functions are only beginning to emerge [2]. Recent reports suggest roles for circRNAs in regulating transcription, protein binding, and sequestration of microRNAs [11–13]. Some circRNAs can also be translated via cap-independent mechanisms to generate proteins [14–16]. Recent work implicates circRNAs in antiviral immunity [17, 18]. Expression patterns of circRNAs in the brain suggest that they might serve important functions in the nervous system [19].

Several RNA-seq studies have found that circRNAs are differentially expressed during aging. Over 250 circRNAs increased in expression within *Drosophila* head tissue 1 to 20 days old [20]. Trends for increased circRNA expression have also been identified during embryonic/postnatal mouse development [10, 21, 22], suggesting that circRNA accumulation might begin early in development. We recently reported that circRNAs were biased for age-accumulation in the mouse brain [23]. In hippocampus and cortex, ∼ 5% of expressed circRNAs were found to increase from 1 month to 22 months of age, whereas ∼1% decreased [23]. This accumulation trend was independent of linear RNA changes from cognate genes and thus was not attributed to transcriptional regulation. CircRNA accumulation during aging might be a result of the enhanced stability of circRNAs compared to linear RNAs [24, 25]. Age-related deregulation of alternative splicing [26, 27] leading to increased circRNA biogenesis might also play a role.

*C. elegans* is a powerful model organism for studying aging. Previously, thousands of circRNAs were annotated from RNA-seq data obtained from *C. elegans* sperm, oocytes, embryos, and unsynchronized young adults [4, 24]. Here, we annotated circRNAs from very deep total RNA-seq data obtained from *C. elegans* at different aging time points, uncovering 575 novel circRNAs. A massive trend for increased circRNA levels with age was identified. This age-accumulation was independent of linear RNA changes from shared host genes. Our findings suggest that circRNA resistance to degradation in post-mitotic cells is largely responsible for the age-upregulation trends identified both here in *C. elegans,* and possibly in neural tissues of other animals.

## Results

### Genomic features of circRNAs in *C. elegans*

We set out to map *C. elegans* circRNAs genome-wide and quantify their expression at different ages using RNA-seq. We performed an aging paradigm of wild-type Bristol N2 worms that involved treatment with 5-fluoro-2-deoxyuridine (FuDR) to prevent egg-hatching. RNA from whole worms from three independent biological replicates corresponding to four aging time-points were collected (three replicates each): L4-larval stage (L4), Day-1 (D-1), Day-7 (D-7), and Day-10 (D-10) (Fig. S1). Ribo-depleted total RNA-seq library preparation was performed, followed by sequencing using paired-end 125 nt reads. *De novo* mapping of circRNAs from these total RNA-seq datasets was performed using the find_circ algorithm [24] (see Methods), and with the added restriction of only annotating circRNAs that shared known exonic splice sites (ce11 UCSC genome). This strategy for mapping circRNAs requires the use of only “back-spliced reads” (Fig. 1A), which represent a very low percentage of a typical RNA-seq run (Table S1). This approach was required to distinguish reads corresponding to circRNAs versus their linear counterparts that share the same exons. Of the ∼1.9 billion paired end reads generated, only 111,895 reads (0.006% of total) mapped to circRNA junctions after removal of PCR duplicate reads (Table S1). To annotate circRNAs with high confidence, we used a cut-off of 12 aligned reads per circRNA across the 12 libraries. This minimum read cut-off was more stringent compared to previous circRNA annotations in *C. elegans* [4, 24]. Using our annotation pipeline (Fig. S2), we confidently identified a total of 1,166 circRNAs. In this high confidence list, 591 circRNAs were previously annotated [4, 24], 575 were novel (Fig. 1B).

**Figure 1.**
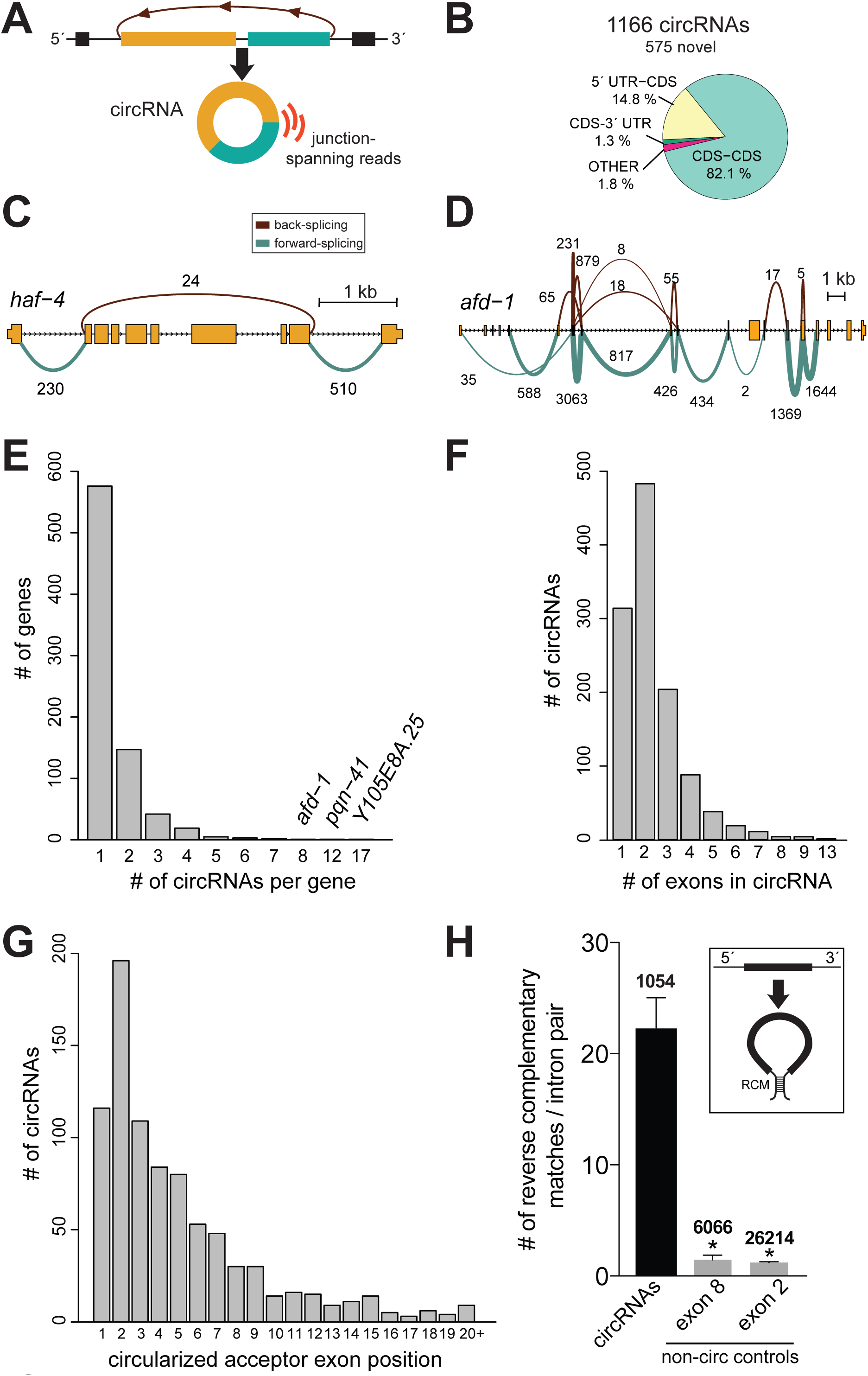
Genomic features of *C. elegans*circRNAs. **A)** Schematic showing a circRNA generated by backsplicing of exons, and the mapping of reads to the back-spliced junction. **B)** Distribution of circRNAs in *C. elegans* genome. Data was mapped from 12 total RNA-seq libraries of N2 worms, including L4 larvae (L4), Day 1 (D-1), Day 7 (D-7), and Day 10 (D-10). CDS, protein coding sequence. **C)** Forward and reverse splicing patterns for the *haf-4* gene. Linear spliced read count (green) and back-splicing read count (brown) are shown. Numbers correspond to the number of spliced reads detected in the D-10 datasets. Only reads corresponding to the junctions included in circRNAs are shown. The gene *haf-4* generates a single circRNA that extends across 8 exons. **D)** *afd-1* generates 8 circRNAs. **E)** Bar plot showing the number of expressed circRNAs per gene. **F)** Number of exons contained within exonic circRNAs. **G)** Ranked position of circRNA first exon for circRNAs containing more than 1 exon. **H)** Presence of Reverse Complementary Matches (RCM) in introns flanking circRNA exons is greater than non-circRNA generating exon controls. Number above bars correspond to # of loci. *, *P* < 0.0001 on Kruskal Wallis test with Dunn’s multiple comparisons.

Most of the 1,166 circRNAs mapped to coding-sequence (CDS) regions of exons (82.1%), followed by circRNAs mapping to exons encompassing 5´ UTR regions and CDS (14.8%) (Fig. 1B). We found that 797 genes express at least one circRNA. As shown for *haf-4* (Fig. 1C), most genes that produced circRNAs expressed a single circRNA (576/797 genes). On the other hand, some genes were found to generate a large number of circRNAs. For instance, the *afd-1* gene was found to generate 8 different circRNAs (Fig. 1D). Overall, 221 out of 797 genes generating two or more circRNAs (Fig. 1E). The number of exons within circRNAs ranged from 1-13, but it was most common for them to harbor 2 exons (Fig. 1F). Only 6.6 % of the 1,166 circRNAs contained 5 or more exons. The reliance of this analysis on back-spliced reads precludes the determination of whether these multi-exon circRNAs have introns removed. As previously found for *Drosophila* and mice [20, 23] there was a bias for circRNAs to emanate from the 5´ end of genes (Fig. 1G).

Base pairing between introns flanking circularizing exons are thought to bring 5´ and 3´ splice sites in close proximity to promote circRNA biogenesis over linear-splicing (Fig. 1H). We used BLAST alignment of introns that flank circRNA-forming exons to identify reverse complementary matches (RCMs). We found that RCMs flanking circRNA loci were strongly enriched (*P* < 0.0001, Kruskal-Wallis test with Dunn’s multiple comparisons test) compared to analogous introns flanking non-circularizing exons (Fig. 1H). Thus, consistent with previous reports [4], our analysis shows that *C. elegans* circRNAs tend to be flanked by introns that pair with one another.

### Experimental validation of circRNAs

We next performed experimental validation of individual circRNAs annotated from our pipeline. One validation method was to prepare cDNA using random hexamers from total RNA, and then perform PCR using outward facing primers that should only amplify a back-spliced circRNA (Fig. 2A). The presence of a back-spliced junction was confirmed for 10/10 circRNAs tested by Sanger sequencing of RT-PCR products (Table S2). In addition, we confirmed a subset of circRNAs by treating total RNA with the exoribonuclease, RNase R, which is known to preferentially degrade linear RNAs over circRNAs [5, 24]. RT-qPCR experiments show that linear RNA *cdc-42* was susceptible to degradation by RNase R, whereas 4/4 circRNAs tested were enriched upon RNase R treatment (Fig. 2B).

**Figure 2.**
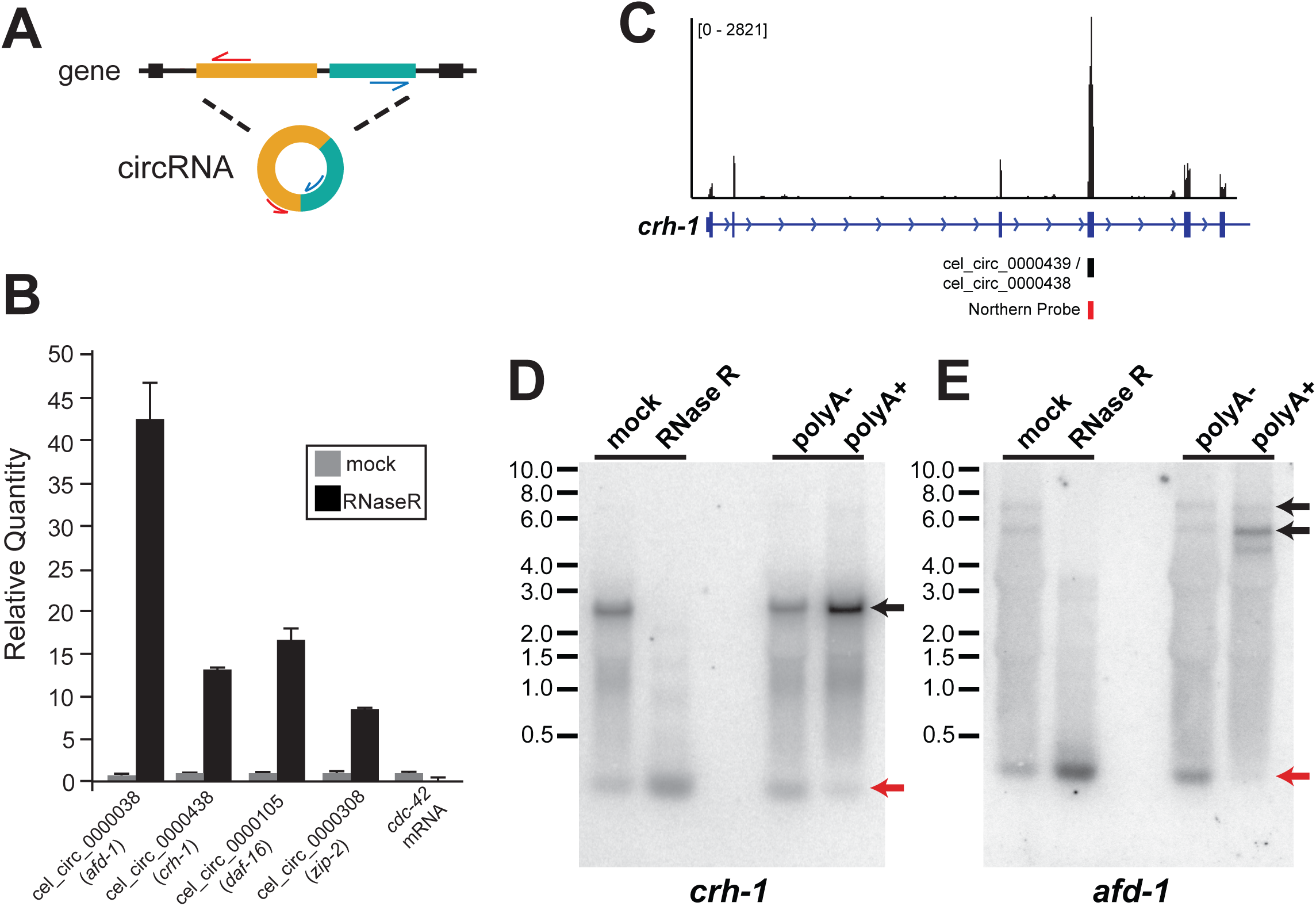
Experimental validation of circRNAs. **A)** RT-PCR strategy to detect circRNAs exclusively using outward facing primer sets. Sanger sequencing of PCR products confirmed 10/10 circRNAs tested (Table S2). **B)** RT-qPCR experiments on RNase R treated mixed age adult worms. Equal amounts of mock-treated and RNase R treated RNA were used for cDNA preparation prior to qPCR. Note the enrichment of circRNAs with RNase R treatment, whereas linear *cdc-42* mRNA is not enriched. **C)** RNA-seq track visualized using Integrated Genomics Viewer from D-7 worms showing read pileup at the *crh-1* gene. Note the increased read number overlapping the circularized exon. cel_circ_0000438 and cel_circ_0000439 differ by 9 nucleotides in length at the 5´ end of the exon. **D)** Northern blot using a probe overlapping the circularized exon of *crh-1* (see panel C) detects bands corresponding to circRNA and mRNA from mixed age adult RNA. Relative circRNA to mRNA abundance is enriched in RNase R treated and PolyA-samples compared to polyA+ samples. Red arrows denote circRNA bands. Black arrows denote likely linear RNA bands. **E)** Northern blot performed using a probe that detects *afd-1* circRNA and mRNA.

Although most circRNAs were of lower abundance compared to linear RNAs from the same gene, some circRNAs annotated were of relatively high abundance. We set out to confirm two of these high abundance circRNAs using Northern blot analysis. In the case of the *crh-1* gene, an increased abundance of reads aligning to an exon harboring a circRNA was clearly evident from visualization of Integrated Genomics Viewer tracks (Fig. 2C). We performed Northern analysis using a probe targeting this circularized exon. This probe should detect both linear and circular transcripts of the *crh-1* gene. As expected, an abundant circRNA migrating at the predicted molecular weight was detected (Fig. 2D). The expression of higher molecular weight linear RNAs was found to be diminished by RNase R treatment, whereas the circRNA bands were unaffected by RNase R treatment (Fig. 2D). We prepared polyA + RNA from a column-based preparation, and collected and precipitated the unselected RNA (polyA-depleted). We found that polyA+ RNA had depleted circRNA levels relative to linear RNA. In contrast, polyA-samples showed enhanced levels of circRNA relative to linear RNA (Fig. 2D). Analogous results were obtained using a probe for an abundant circRNA from the *afd-1* gene (Fig. 2E). Together, these validations provide experimental support that our annotation pipeline detected bonafide circRNAs in *C. elegans*.

### Global circRNA levels dramatically increase during aging

We next quantified the abundance of circRNAs from the different aging time-points. CircRNA read counts were normalized to their corresponding library size to obtain Transcripts Per Million reads (TPM) (Table S3). Principal Component Analysis (PCA) on the circRNA TPM values was performed on the 12 RNA-seq libraries (Fig. 3A). Strikingly, a close clustering of L4 to D-1, and of D-7 to D-10 was observed, suggesting that global circRNA expression levels reflect age.

**Figure 3.**
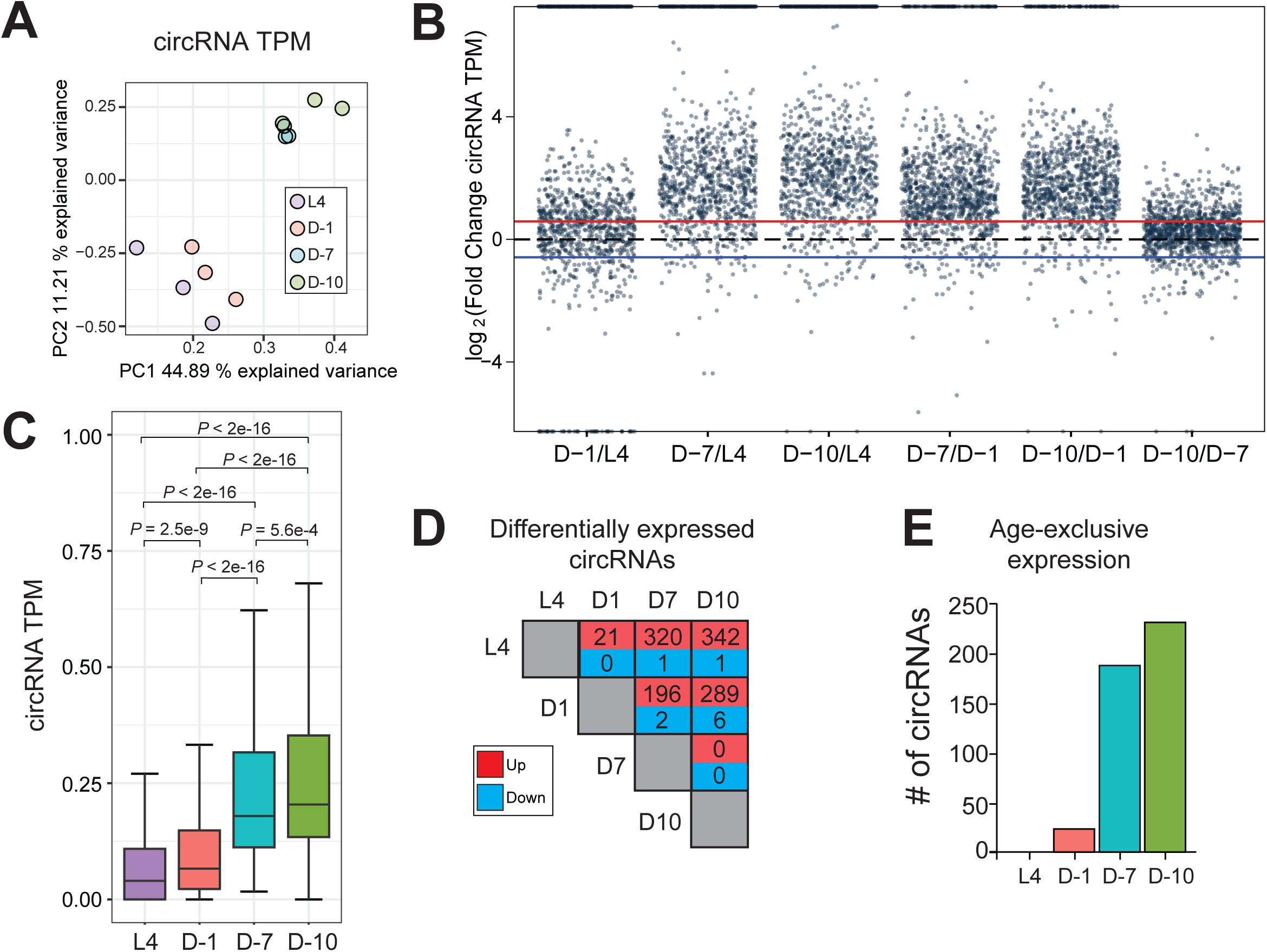
Global circRNA accumulation during aging. **A)** Principal component analysis (PCA) of circRNA Transcripts Per Million reads (TPM) shows clear clustering of young (L4, D-1) versus old ages (D-7, D-10). **B)** Plot of circRNA TPM fold changes in aging time-point pairwise comparisons. Red line represents 1.5-fold increase and blue line represents 1.5-fold decrease. **C)** CircRNA TPM compared among the four aging time-points: L4 larvae (L4), Day 1 (D-1), Day 7 (D-7) and Day 10 (D-10). *P* values reflect non-parametrical Kruskal-Wallis with Nemenyi post-hoc test for multiple comparisons. **D)** Pairwise comparisons of age-increased and decreased circRNAs among the aging time-points (>1.5 FC, *P* < 0.05, FDR < 0.2). **E)** Histogram showing number of circRNAs exclusively expressed at a single time-point (> 3 reads among 3 biological replicates).

To further investigate trends in circRNA levels during aging, we plotted circRNA log_2_ fold changes in TPM for all pairwise comparisons of the aging time-points (Fig. 3B). We found that 1,052 circRNAs (90.2%) were at least 1.5-fold greater in D-10 versus L4 time-points, whereas only 37 circRNAs were 1.5-fold greater in L4. Similar trends were found in other pairwise comparisons between older (D-10, L4) versus younger (D-1, L4) time-points (Fig. 3B). For instance, in comparisons between D-7 versus D-1, 80.8% of circRNAs were >1.5-fold higher in the older time-point.

To gain statistical support for these dramatic aging trends we performed several additional analyses. The global expression of circRNA TPM values was compared across ages by non-parametrical Kruskal Wallis test with Nemenyi post-hoc test for multiple comparisons (Fig. 3C). Interestingly, comparisons between more distant aging time-points (D-10/L4, D-7/L4, D-10/D-1, and D-7/D-1) yielded the lowest *P* values (*P* < 2E-16). In contrast, the D-1/L4 and D-10/D-7 comparisons had less significant *P* values (*P* = 2.5E-9 and 5.6E-4, respectively), which might reflect the ages being closer together.

In order to identify the individual circRNAs with statistically significant changes in expression during *C. elegans* aging, we performed *t*-tests on TPM values between each aging time-point (*P* < 0.05, > 1.5 Fold Change (FC), False Discovery Rate (FDR) < 0.2). An overwhelming bias for upregulation of circRNAs during aging was uncovered. For instance, in the comparison of D-7 versus D-1, a total of 196 circRNAs were upregulated whereas only 2 were downregulated (Fig. 3D). Comparing D-10 vs L4 age time-points, 342 circRNAs were upregulated, whereas only 1 was downregulated (Fig. 3D). This aging trend was also observed when we lowered our stringent expression cutoffs to include low expressed circRNAs by reducing the minimum number of reads required across libraries to 6. Setting a minimum level of 3 reads per time-point as the cutoff, we plotted time-point specific circRNAs. We found that D-7 and D-10 clearly expressed the greatest number of time-point specific circRNAs (Fig. 3E). In fact, there were no circRNAs exclusively expressed in L4 worms and not other aging time-points (Fig. 3E). Together, these data demonstrate that circRNAs show an overwhelming bias for age-accumulation in *C. elegans*.

### Experimental validation of circRNA age accumulation

We next set out to confirm circRNA expression trends for particular circRNAs that are generated from genes with interesting functions. We performed RT-qPCR validation for 9 individual circRNAs, and included circRNAs generated from genes that are involved in lifespan determination (*akt-1*, *crh-1*, *daf-16*, *daf-2*). We selected circRNAs with a variable range of overall expression levels for validation (Fig. 4A). Of these 9 circRNAs, 7 met the stringent FDR value for differential expression (increased) between at least one old versus young time-point. Three of these circRNAs did not meet statistical significance for differential expression (D-10 versus L4) from the RNA-seq data, including two circRNAs of low abundance (Fig. 4A). Quantification of these same circRNAs by RT-qPCR revealed that 9/9 circRNAs tested were significantly upregulated in D-10 vs L4 (Fig. 4B). RT-qPCR analysis also showed that the tested circRNAs did not continue to increase in D-10 animals, consistent with the RNA-seq results (Fig. 4B).Notably, RT-qPCR validation for most circRNAs showed fold-changes greater than those detected by RNA-seq differential expression analysis. Most of these qPCR quantified changes were >10-fold between L4 and D-7. Remarkably, changes in circRNAs from the *gld-2* and *daf-16* loci were >40-fold induced between L4 and D-7 (Fig. 4B). Overall, these qRT-PCR validations strongly support the trend of greater circRNA abundance in old (D-7, D-10) versus young (L4, D-1) animals. These confirmations also suggest that the actual number of circRNAs that increase during aging is much greater than what was found to significantly change from the RNA-seq analysis (Fig. 3D).

**Figure 4.**
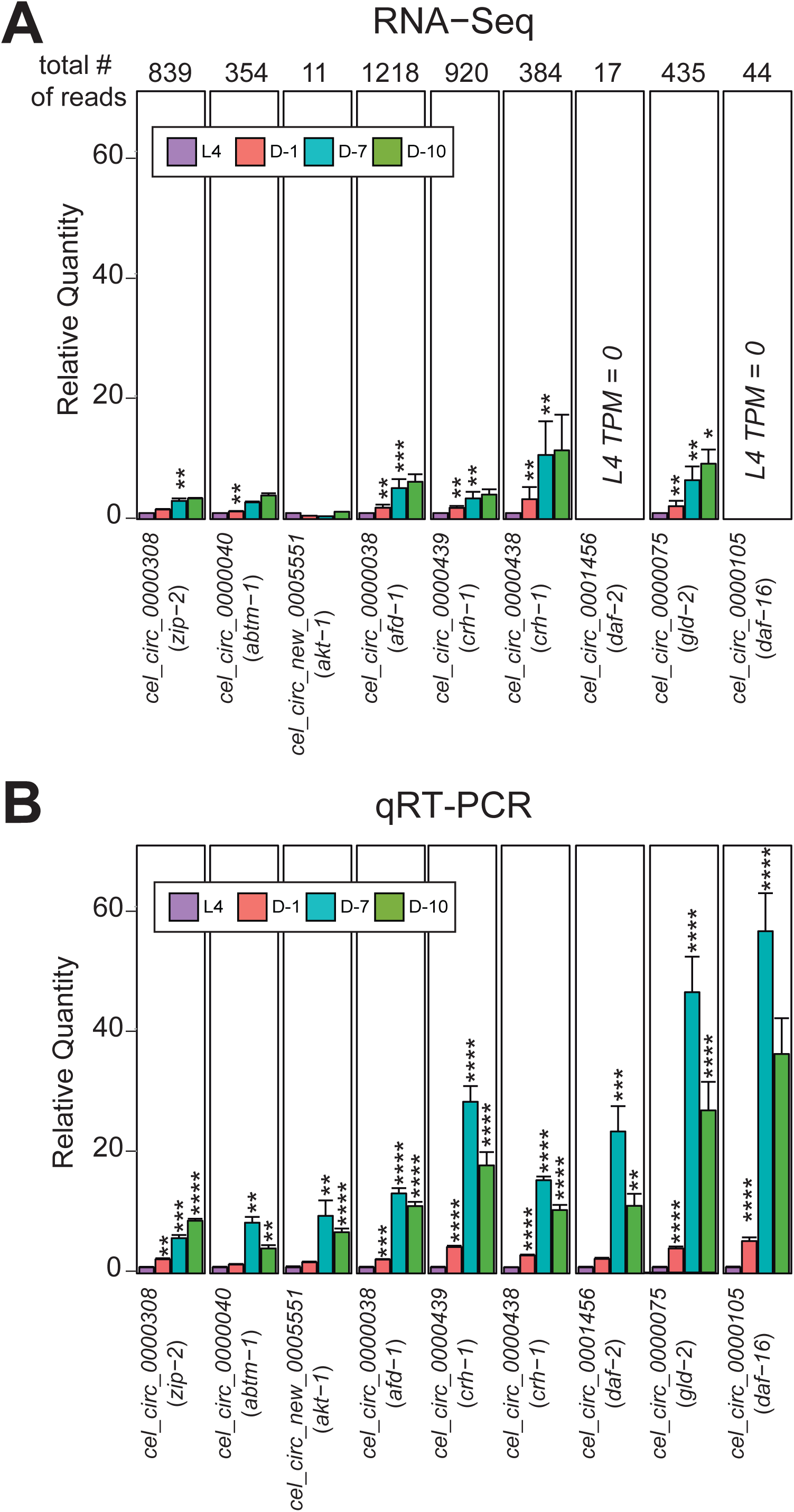
Validation of circRNA age-accumulation. **A)** RNA-seq quantification of select circRNAs during aging (TPM fold-change with L4 set at 1). Total # of reads across all libraries for each circRNA is noted above graph. Labels display circRNA names with host gene in brackets. **B)** qRT-PCR data for the same selected circRNAs as in B). Data is normalized to *cdc42* mRNA. Note the greater magnitude of age-related expression changes reported by qRT-PCR versus RNA-seq for all circRNAs. Error bars represent SEM. *, *P* ≤ 0.05; **, *P* ≤ 0.01; ***, *P* ≤ 0.001; ****, *P* ≤ 0.0001.

### CircRNAs show greater age-related increase than linear RNAs

Next, we performed differential expression analysis on linear RNAs among the different age time-points. Linear RNAs previously found to be differentially regulated during aging displayed similar expression trends in our datasets. For example, between D-7 and D-1 *hsp-70* and *cht-1* were upregulated during aging, whereas *fat-7*, *ifp-1*, and *ifd-1* were downregulated (Table S4). In contrast to circRNA trends, a global bias for linear RNA differential expression was not evident. Similar numbers of upregulated and downregulated linear RNAs between aging time points were identified using CuffDiff (Table S4). Scatterplots comparing old versus young timepoints for linear RNA levels (Fig. 5A-D), and circRNA levels (Fig. 5E-H) exemplify the stark contrast in the age-related trends. Across all the old versus young datasets, it is clear that nearly all differentially expressed circRNAs are increased during aging, whereas similar numbers of linear RNAs are increased and decreased.

**Figure 5.**
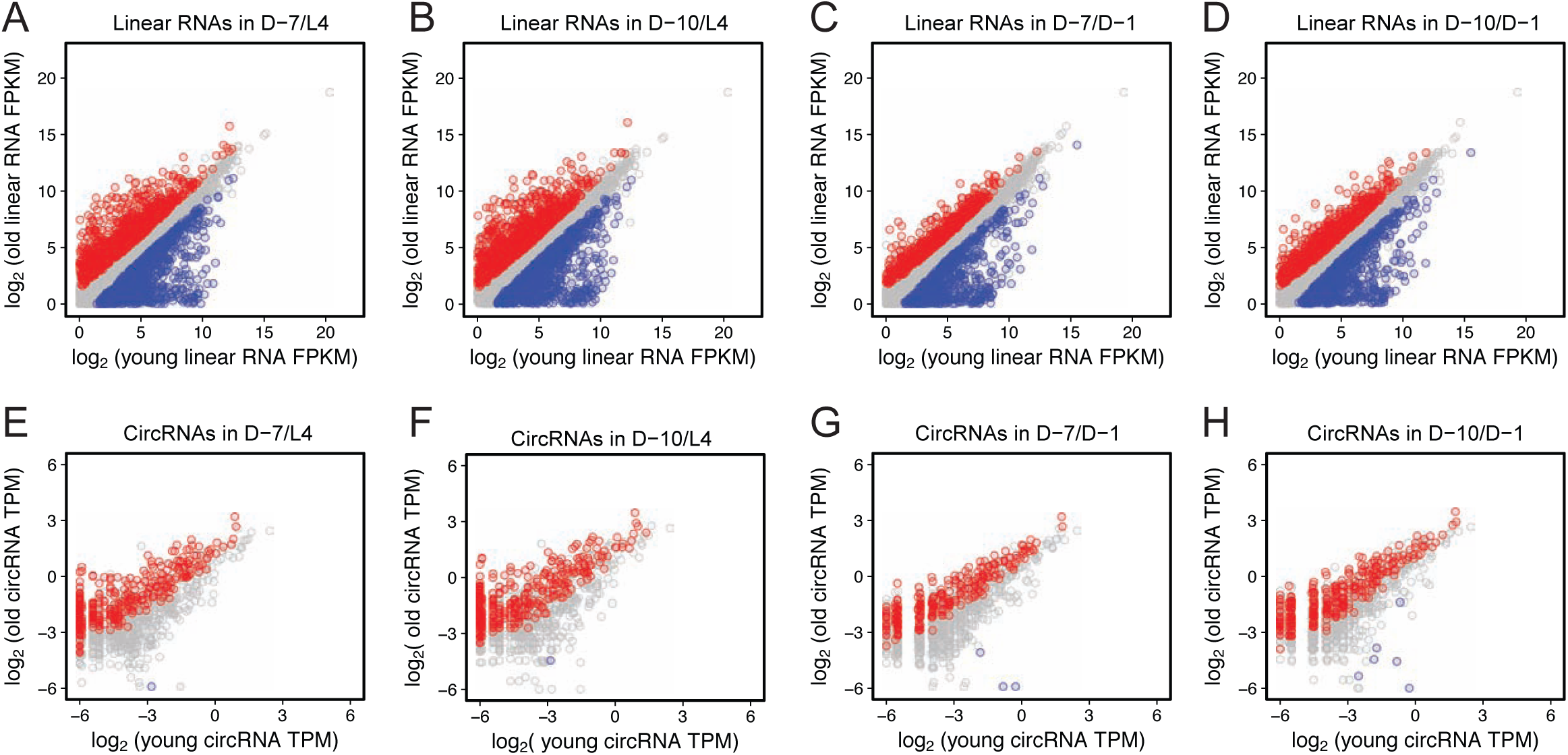
Comparison of circRNA and linear RNA expression during aging. **A)** Scatterplot showing pairwise comparisons between aging time-points for linear RNA levels (**A-D**) and circRNAs (**E-H**). Log_2_ linear RNA FPKM value scatterplots are shown for **A)** D-7 vs L4, **B)** D-10 vs L4, **C)** D-7 vs D-1, and **D)** D-10 vs D-1. Log_2_ circRNA TPM scatterplots are shown for **E)** D-7 vs L4, **F)** D-10 vs L4, **G)** D-7 vs D-1, and **H)** D-10 vs D-1. Significant changes have a fold-change > 1.5, *P* < 0.05, FDR < 0.2. Red data points show age-upregulated circRNAs, whereas blue data points show downregulated circRNAs.

Although linear RNAs lacked a global bias for increased levels during aging, it was still possible that increased transcription of circRNA-hosting genes could contribute to the circRNA expression trends. Thus, we analyzed whether circRNA accumulation was independent of host-gene expression. Density plots were generated to contrast circRNA fold-changes versus their counterpart linear RNA changes from the same host gene (Fig. 6). For this analysis, we used an expanded list of circRNAs, requiring at least 3 reads per age time-point. In the expected old versus young time-point comparisons, a clear upward vertical shift was evident in the density plots (reflecting increased circRNA expression), and only a minor horizontal shift to the right (reflecting increased linear RNA expression) (Figs. 6B, 6C, 6D, 6E). This suggests that circRNA accumulation trends are largely independent of linear RNA changes. For comparisons between closer time points (D-1/L4 and D-10/D-7) the density plots lacked clear vertical or horizontal shifts (Figs. 6A, 6F). Thus, circRNAs globally accumulate during aging in *C. elegans* independently of general changes in expression from their host genes.

**Figure 6.**
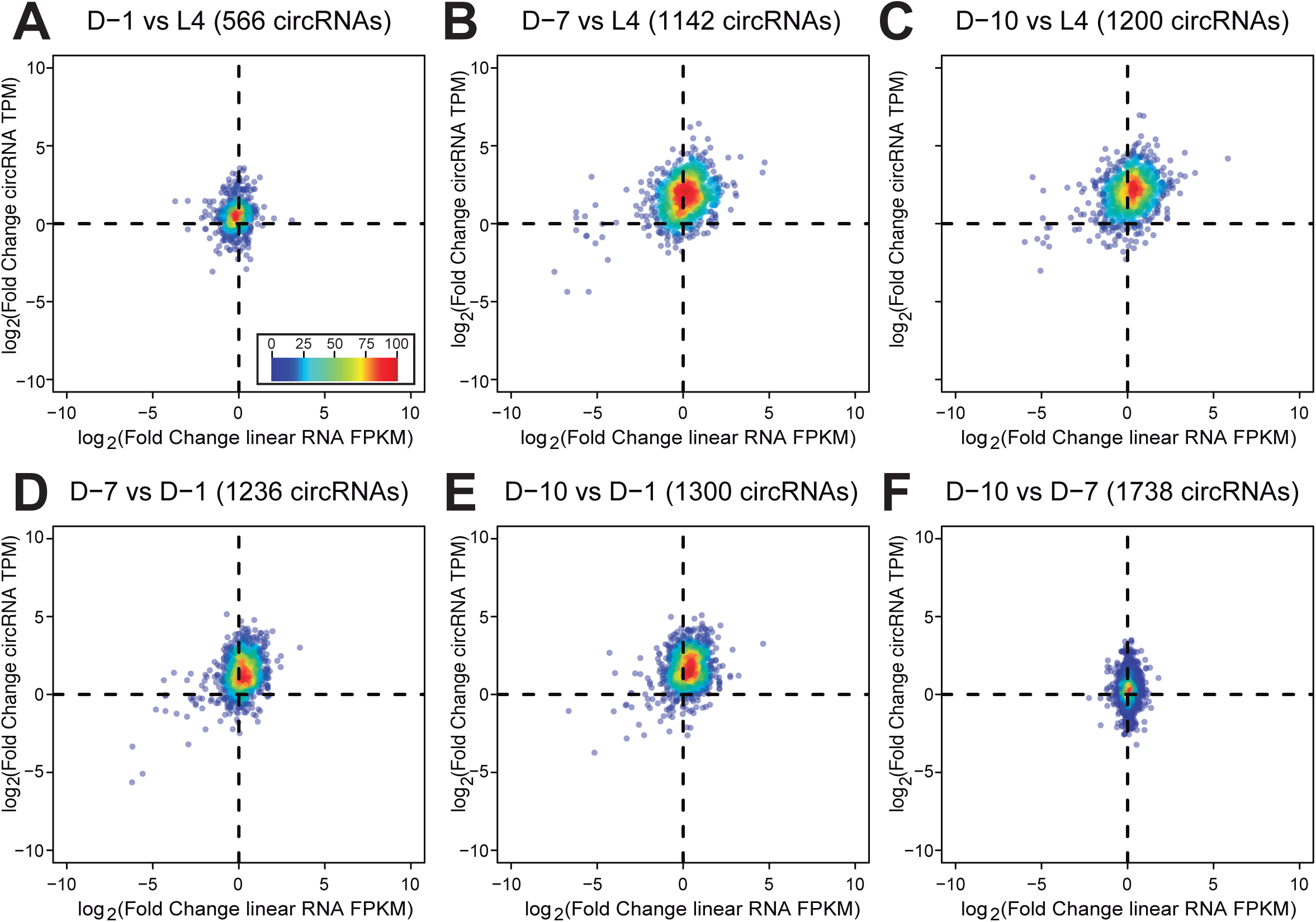
Age accumulation of circRNAs is independent of host gene expression. Density plots for circRNA TPM fold-change versus linear RNA FPKM fold-change. Log_2_ fold changes of circRNAs versus log_2_ fold changes of linear RNAs from parental genes are shown. **A)** D-1 vs L4, **B)** D-7 vs L4, **C)** D-10 vs L4, **D)** D-7 vs D-1, and **E)** D-10 vs D-1, **F)** D-10 vs D-7. Scale bar inset in panel A represents circRNA number and applies to all the density plots. For old versus young time-point comparisons, it is evident that upregulation of circRNAs is largely independent of linear RNA expression from the same gene (upward shift in plots). Plots include circRNAs with > 3 reads for each time-point under comparison.

### Discussion

This study is the first to report an age-associated accumulation of circRNAs in *C. elegans*. Previous studies have documented the bias for circRNAs to be increased during aging in neural tissues of *Drosophila* and mice [20, 23]. Interestingly, the trends uncovered here during *C. elegans* aging are much more dramatic. Many confirmed expression trends were >10 fold changed between L4 and D-7. Of the hundreds of differentially expressed circRNAs, the vast majority increased with age (Figs. 3, 4, 5). Experimental confirmations of the age-accumulation trends by qRT-PCR were of greater magnitude than those reported from the RNA-seq data, and qRT-PCR experiments revealed several circRNAs to accumulate with age that were not significantly increased in the RNA-seq data (Fig. 4). Given this evidence, we surmise that most circRNAs expressed in *C. elegans* accumulate with age, and that many of those not passing statistical thresholds for differential expression are simply of too low abundance to be quantified by the limited number of back-spliced RNA-seq reads in these datasets.

Why is the age-accumulation trend of circRNAs in *C. elegans* much stronger than in other organisms tested so far? After completing development and a brief reproductive period, *C. elegans* spends the remainder of its adult life comprised almost exclusively of post-mitotic cells [28]. The FuDR treatment employed in this study inhibits DNA synthesis and is commonly used to prevent egg-hatching in aging and life-span studies of *C. elegans* [29] and also reduces the presence of proliferating cells. We have previously proposed that either the high stability of circRNAs or alterations in alternative splicing with age could explain age-upregulation circRNA trends [2]. The trends observed here in *C. elegans* argue strongly for a low rate of circRNA decay being the responsible mechanism. We propose that the dramatic genome-wide increase of circRNA levels during aging are a consequence of the dominance of post-mitotic cells in *C. elegans* adult worms combined with the high stability of circRNAs. As neurons are post-mitotic, perhaps this can also explain why age accumulation is most notable in *Drosophila* heads (which are rich in neurons) [20] and in brain regions of mice [23].

To investigate the potential functional significance of age-accumulated circRNAs, we performed Gene Ontology analysis on the host genes of expressed *C. elegans* circRNAs, and found many significantly enriched categories, including an enrichment in the Biological Process category of “determination of adult lifespan” (Fig. S3, Table S5). A clear bias for particular cellular components, functions, or biological roles was, however, not uncovered by these efforts. It is of course possible that *trans* functions of circRNAs are completely distinct from the curated roles of their host genes.

Combined with previous studies in *Drosophila* [20] and mice [23], we here provide further evidence that circRNA age-accumulation is a broadly conserved pattern. Future work can now take advantage of the powerful genetics of *C. elegans* to delineate aging functions of individual circRNAs. Generating loss-of-circRNA mutants in *C. elegans* by disrupting base pairing of flanking introns could be a fruitful approach. Various RNAs found to be differentially regulated during aging were subsequently found to impact lifespan in *C. elegans* mutant analysis, including linear RNAs [30], microRNAs [31] and lncRNAs [32]. Given that these mutant studies on lifespan were based on comparatively modest fold-changes during aging, the massive upregulation trends for circRNAs provide solid rationale for disrupting or overexpressing circRNAs in *C. elegans* and testing for effects on lifespan and healthspan. However, one should also consider that the aging process might be impacted generally by the total compendium of hundreds of circRNAs accumulating in cells, as opposed to individual circRNAs. Thus, non-conventional approaches to alter the expression of many circRNAs simultaneously might be required to uncover age-related functions of circRNAs.

## Experimental Procedures

### *C. elegans* maintenance and culturing

The *C. elegans* Bristol N2 wild type strain was grown and maintained as previously described [33]. To synchronize populations, gravid adults were bleached, eggs were collected and left overnight in 1X M9 buffer with rocking. Starvation arrested L1 larvae were placed on 150x100mm NGM plates with 10X concentrated *E. coli* OP50 as a primary food source, and kept at 15°C. At the L4 larval stage, animals were collected using a 25 µm nylon mesh (Sefar) and either harvested for RNA extraction, or placed on 150x100mm NGM plates containing 75 mM 5-fluoro-2´-deoxyuridine (Sigma Aldrich) with 10X concentrated *E. coli* OP50 and kept at 15°C until animals were harvested at Day-1, Day-7 and Day-10 of adulthood (see Fig. S1).

### RNA extraction and library preparation

A 250 µl mixture of animals in 1X M9 buffer was added to 750 µl of TRIzol LS (ThermoFisher) and immediately frozen with liquid N_2_. Lysates were freeze/thawed at −80°C, disrupted with Mixer Mill 400 (Retsch) and dounce homogenizer (Corning) to break apart the cuticle of animals. Any cellular debris was removed by low-speed centrifugation. RNA was extracted using the Purelink RNA mini-kit with DNAse I treatment (Ambion). RNA quality was assessed by Bioanalyzer (Agilent) and quantified using Quant-iT RiboGreen RNA Assay kit (ThermoFisher Scientific).

### Library preparation and high-throughput sequencing

Libraries were prepared using the Illumina TruSeq Stranded Total RNA Library Prep Kit as recommended by the manufacturer (Illumina) with modified conditions to increase the size of the cloned fragments (fragmentation at 85°C x 5 min). Barcoded libraries were sequenced at New York Genome Center (New York, NY) using Illumina HiSeq 2500 system to obtain paired-end 125 nt reads. Raw FASTQ files from the RNA-seq data were deposited at the NCBI Sequence Read Archive (accession numbers are listed in Table S1).

### Experimental Validation of circRNAs

To confirm individual circRNAs, RNA was reverse transcribed using Superscript III with random hexamers (Invitrogen). PCR products were gel extracted then Sanger sequenced, or first cloned into the PCR 2.1-TOPO TA vector (Invitrogen) prior to Sanger sequencing (Nevada Genomics Center, University of Nevada, Reno). For qPCR analysis, we used a BioRad CFX96 real time PCR machine with SYBR select mastermix for CFX (Applied Biosystems) using the delta delta Ct method for quantification. These experiments were performed using technical quadruplicates. Total RNA from *C. elegans* was treated with or without 0.4 U/µL RNase R (Epicentre) with 2U/µL RNaseOUT (ThermoFisher Scientific) for 2 hours at 37°C. RNase R reactions were terminated with 0.5% SDS buffer (0.5% SDS, 10 mM Tris–HCl [pH 7.5], 1.25 mM EDTA [pH 8], 100 mM NaCl) and RNA was purified from reactions using acid phenol chloroform extraction with isopropanol precipitation and 70% ethanol washes. Equal amounts of RNase R or mock treated RNA served as input for cDNA preparation.

PolyA+ RNA and polyA-RNA were obtained from total RNA using NucleoTrap mRNA kit (Machery-Nagel). RNA bound to oligo(dT) beads was carried through the complete polyA+ enrichment according to manufacturer’s protocol while RNA that remained in the supernatant (unbound to the oligo(dT) beads) was precipitated with isopropanol and washed with 70% ethanol. Northern analysis was performed as previously described [23], with probe hybridization taking place overnight at 42°C, and all blot washing steps at 50°C.

### CircRNA prediction and mapping

For *de novo* identification of circRNAs, a computational pipeline was carried out as previously described with filtering for duplicate reads and removing circRNA annotations spanning multiple genes [23] (Fig. S2). We obtained circRNA junction spanning FASTA sequence templates of 200 nucleotides using the *C. elegans* genome (from WBcel235 ce11 UCSC genome) as a reference. Assignment of circRNAs to their corresponding parental genes was performed using custom R scripts based on the library GenomicFeatures [34].

### circRNA normalization and cutoffs

A minimum of 12 reads across all the libraries was required for each circRNA in order to be considered for downstream analyses. To account for variability due to differences in library size, counts attributed to individual circRNAs were normalized to Transcripts Per Million reads (TPM). We calculated the corresponding fold changes by pair-wise comparisons of the average TPMs between time points. Individual unpaired *t*-tests were performed across the correspondent TPM values by pair conditions. *P* values were corrected for multiple hypothesis testing with False Discovery rate (FDR) < 0.2. To identify stage-specific changes (Fig. 3E, Fig. 6), we required a minimum of 3 reads per time-point.

### circRNA expression and plots

ggplot2 (Wickham, 2012) and ggrepel (https://cran.r-project.org/web/packages/ggrepel/index.html) R libraries were used for scatterplots. Gene models diagrams were generated using the Gviz package [35]. Density plots were generated with the LSD package (https://cran.r-project.org/web/packages/LSD/index.html).

### Quantification of linear expression

Linear RNA-Seq reads were mapped to *C. elegans* ce11 annotation using TopHat [36] with default settings. To quantify the differential expression across time points we used Cuffdiff [36]. Genes with fold changes > 1.5 and a Benjamini-Hochberg corrected *P* value < 0.05 (default parameter used in the Cuffdiff algorithm) were considered to be differentially expressed.

### Gene Ontology analysis

We used the Cytoscape plugin ClueGO [37] (together with the Gene Ontology Annotation (GOA EMBL-EBI) (http://www.ebi.ac.uk/GOA) released on 11/17/2016. Network specificity was set to “medium”. The enrichment statistic used was a two-side hypergeometric test, correcting for multiple testing with the Bonferroni method. The cutoff for considering a term as enriched was set at *P* < 0.05. To reduce the number of redundant terms we used the GO term grouping option which uses a Kappa score to collapse terms that share elements. We set the minimal number of elements in a group at 3. Bar graphs of significant GO terms were created using the enrichment score or – log(*P*-value).

### RCM and motif analysis

To identify Reverse-Complementary Matches (RCMs), pairs of intron sequences flanking the circRNAs were extracted from the *C. elegans* genome (WBcel235, ce11) using custom scripts available at: https://github.com/alexandruioanvoda/IntronPicker. The corresponding sequences were used as input for the RCM analysis using custom scripts (https://github.com/alexandruioanvoda/autoBLAST) that employed BLAST (parameters: blastn, word size 7, output format 5) to identify matches. Exons 2 and 7 from non-circRNA generating exons were used as controls to account for the possibility that intron pairing might be influenced by exon location within genes.

## Acknowledgements

This work was supported by the National Institute of General Medical Sciences grant P20 GM103650 to P.M. and National Institute on Aging grant R15 AG052931 to P.M. The work was also supported by a mICRo grant from the University of Nevada, Reno office of the Vice President for Research and Innovation to A.V.D.L. and P.M. The N2 strain was provided by the CGC, which is funded by NIH Office of Research Infrastructure Programs (P40 OD010440).

## Author contributions

M.C.L, M.G., A.V.D.L. and P.M. wrote the paper. M.G., M.C.L., H.N.G, P.M., and A.V.D.L. conceived the project. M.G., H.N.G., and D.A.C., performed experiments. M.C.L and A.I.V. performed computational analysis of the data. M.C.L, M.G., A.V.D.L., and P.M. analyzed the data.

## Supporting Information

### Supplementary Figure Legends

**Supplementary Figure 1:**
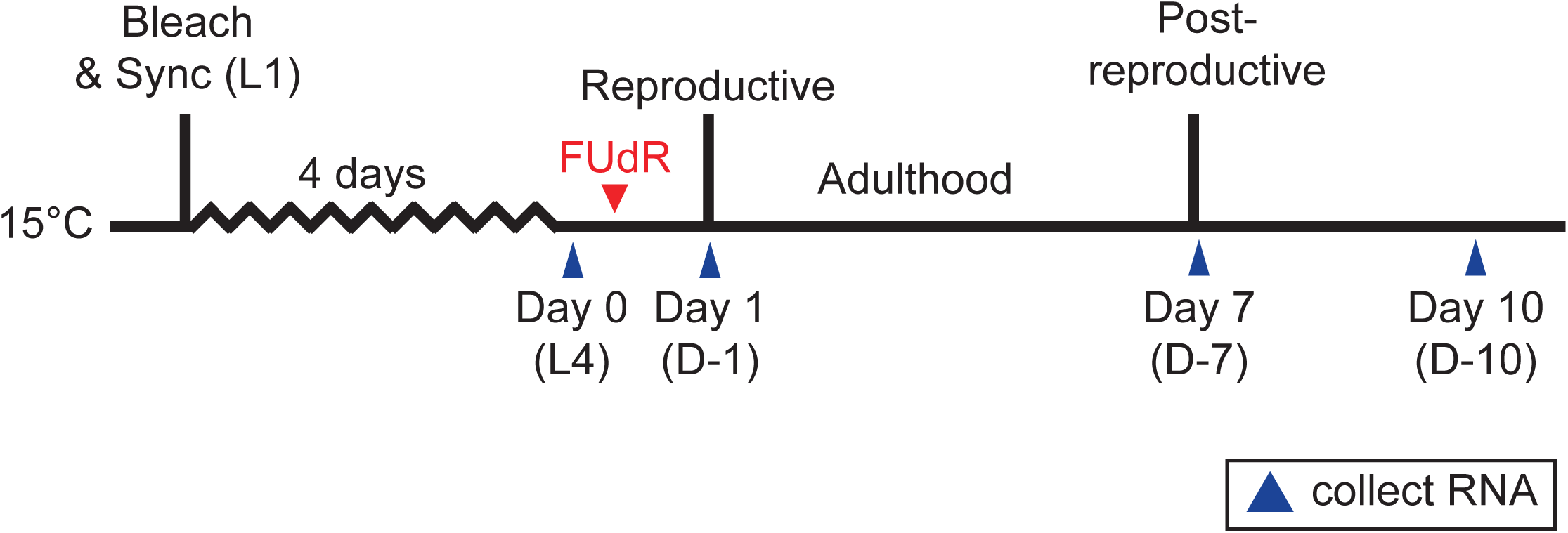
*C. elegans* aging paradigm.

Protocol for collecting total RNA during *C. elegans* aging. Wild-type animals were fed *E. coli* OP50 and grown at 15°C. Gravid adults were bleached and populations were synchronized as L1 larvae and grown for an additional 4 days. At the L4 larval stage (Day 0), animals were either collected or transferred to FUDR containing NGM agar plates seeded with *E. coli* OP50, and were allowed to continue growth at 15°C. Total RNA was collected at different age time-points (L4, D-1, D-7, and D-10).

**Supplementary Figure 2:**
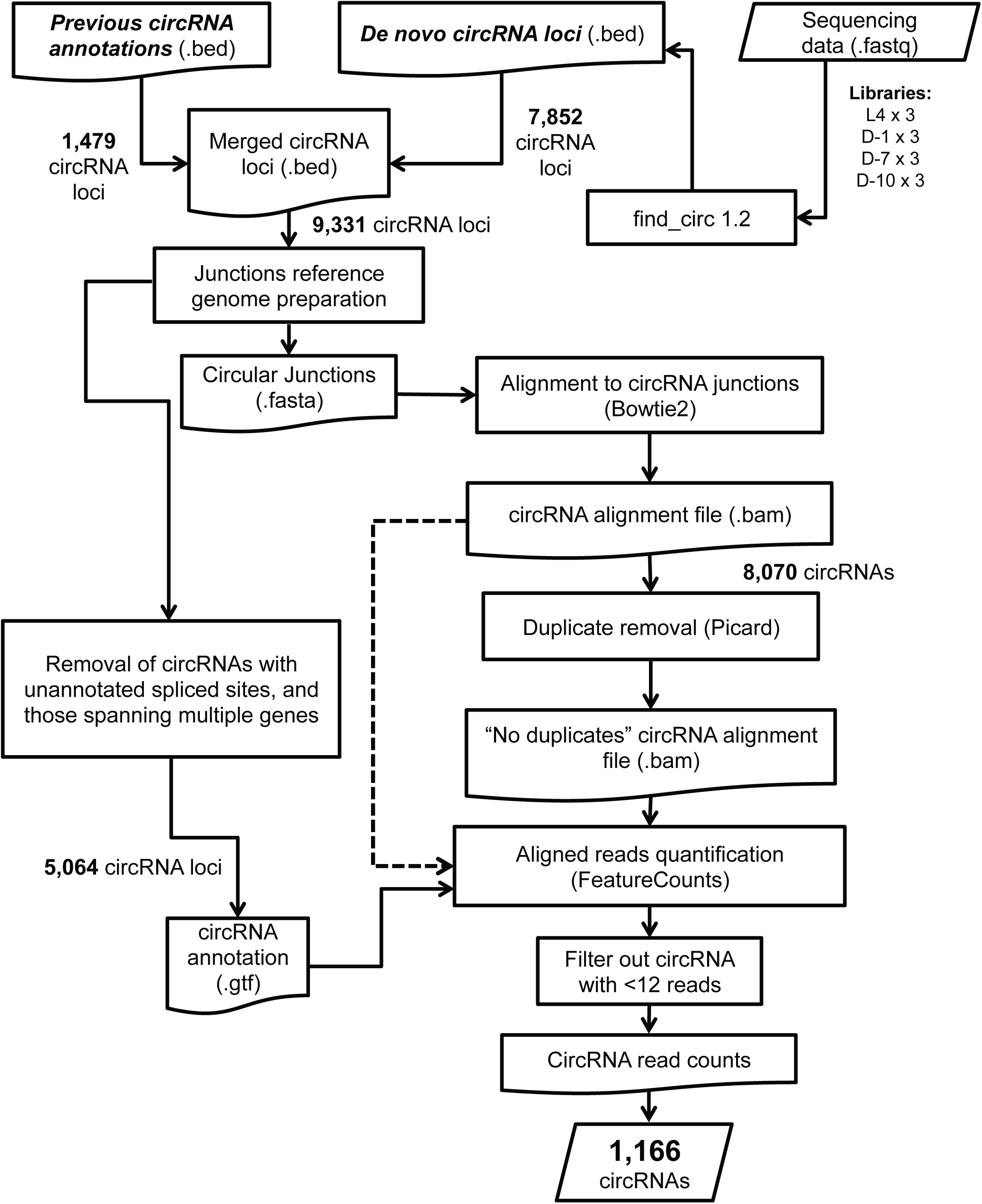
Pipeline for circRNA annotation. A flowchart of the computational pipeline used for circRNA identification.

**Supplementary Figure 3:**
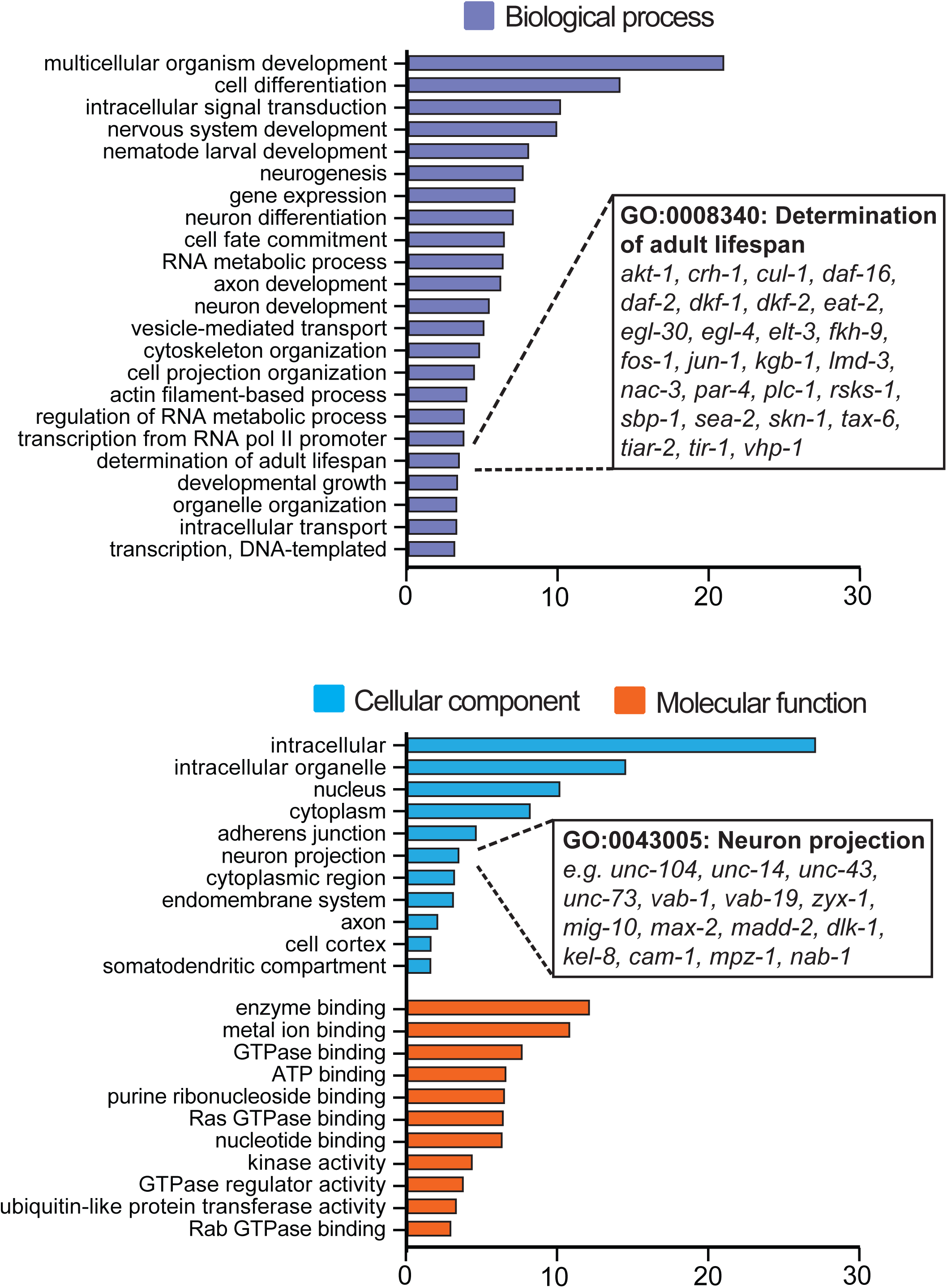
Gene Ontology (GO) analysis of circRNA host genes. Visualization of ClueGO analysis of the 797 genes harboring circRNAs. Complete table of GO analysis is found in Table S5.

### Supplementary Table list

**Table S1:** RNA-seq read statistics

**Table S2**: Oligonucleotides used for experimental validation

**Table S3:** circRNA expression data

**Table S4:** Linear RNA differential expression analysis

**Table S5**: Gene Ontology analysis for circRNA host genes

